# Maintenance of quantitative genetic variation in regions of restricted recombination via associative overdominance

**DOI:** 10.64898/2026.09.20.752755

**Authors:** Yan Zhao, Chaebin Lee, Parul Johri, Brian Charlesworth, Tom R. Booker

**Affiliations:** Department of Forest and Conservation Science, University of British Columbia, Canada; Department of Biology, University of North Carolina at Chapel Hill, USA; Institute of Evolutionary Biology, University of Edinburgh, UK

## Abstract

Mutation-selection balance is the most plausible general mechanism for the maintenance of additive genetic variance (*V*_*A*_) for quantitative traits in natural populations. It is well established that the genetic variants affecting traits (QTLs) are widely distributed across species’ genomes. At the same time, unconditionally deleterious mutations are also expected to be common, with those with the largest fitness effects tending to be strongly recessive. Many genomes possess regions of restricted recombination that could result in tight linkage between largely recessive deleterious alleles, conditions which can lead to elevated heterozygosity via associative overdominance (AOD). Here we show that weakly selected QTLs linked to genomic regions exhibiting AOD can elevate *V*_*A*_ above what is expected in the absence of unconditionally deleterious mutations. Selection on traits when *V*_*A*_ is maintained by AOD, however, may expose recessive deleterious alleles as homozygotes, reducing fitness. Thus, while AOD may contribute to the maintenance of *V*_*A*_, variation maintained in this way may not be readily accessible by natural or artificial selection.

---

The additive genetic variance (*V*_*A*_) of a quantitative trait shapes the paths that its evolution can take, and determines the response to directional selection on a trait jointly with the intensity of selection (1). Substantial *V*_*A*_ is routinely found for quantitative traits in both small and large populations, but genetic studies have struggled to adequately explain its magnitude (2, 3). Observations on continuous phenotypic variation in nature suggest that stabilising selection is common (4). Indeed, the distribution of genetic variation underlying complex traits in humans is consistent with stabilising selection (5). However, stabilising selection on its own tends to erode variation rather than maintain it, leading researchers to focus on models of mutation-selection balance (6).

Surveys of natural populations have typically shown that quantitative traits are generally controlled by numerous loci with small phenotypic effects, in agreement with Fisher’s “infinitesimal model” (7). Population genomic and experimental studies in a wide assortment of species broadly suggest that the distribution of fitness effects (DFE) of new mutations is highly leptokurtic (i.e. it is peaked close to zero with a long tail of deleterious effects). Simple theoretical models and experimental evidence suggest that small effect mutations will tend to be close to additive in their effects, with larger effect mutations being more recessive, and this is supported by population genetic data on deleterious mutations (8).

Recombination rates are known to vary widely across eukaryotic species’ genomes, with large regions of suppressed recombination reported in many species (9). In regions of restricted recombination, therefore, we may expect alleles at weakly selected quantitative trait loci (QTLs) to be tightly linked to more strongly selected, deleterious variants. The fates of weakly selected and neutral alleles are likely be heavily influenced by selection at linked sites (10). Purifying selection against additive and/or strongly deleterious mutations can reduce variability at linked sites via background selection (BGS) (10). Neutral alleles linked to deleterious mutations that are recessive or partially recessive may exhibit elevated heterozygosity due to associative overdominance (AOD) (10), because mutant homozygotes are eliminated, leaving a larger fraction of heterozygotes at neutral sites than expected under neutrality. When multiple deleterious recessive mutations are tightly linked, genotypes possessing combinations of deleterious alleles in negative linkage disequilibrium may be favored, which can elevate variability at linked neutral sites above background levels, a phenomenon referred to as pseudo-overdominance (POD) (11). Note that POD is special case of AOD and for the rest of this paper we refer solely to AOD (10). Elevated heterozygosity and other evidence consistent with AOD has been reported in large regions of restricted recombination present in both maize and pearl millet genomes, for example (12, 13). The *V*_*A*_ contributed by a locus in a randomly mating population is proportional to its heterozygosity (see Methods). Therefore, if QTLs were present in a region exhibiting AOD, we might expect them to exhibit higher heterozygosity than in more highly recombining regions, resulting in an increase in *V*_*A*_ for quantitative traits.

## Results and Discussion

Using simulations, we found that AOD can indeed cause an increase in *V*_*A*_ for traits under stabilising selection, whereas BGS decreases it (Figure 1). We compared the maintenance of *V*_*A*_ for a trait subject to stabilising selection under two contrasting population genetic models. In the first, only mutations that affected the quantitative trait occurred. In the second, we added unconditionally deleterious mutations, with varying scaled selection coefficients (2*N*_*e*_*s*, where *N*_*e*_ is the effective population size, *s* is the strength of selection against homozygotes, and *h* is the dominance coefficient. We compared the magnitude of *V*_*A*_ under these two models in genomes that either recombined uniformly across their length or that contained a central non-recombining region. When chromosomes did not possess a non-recombining region, unconditionally deleterious mutations tended to slightly decrease *V*_*A*_, regardless of the strength of selection or level of dominance (Figure 1). However, when genomes possessed a non-recombining region, the presence of deleterious mutations with 2*N*_*e*_*s* ≤ 10 and *h* ≤ 0.2 led to increases in *V*_*A*_ above what was observed in QTL-only models, with larger increases observed in genomes possessing a larger non-recombining region (Figure 1). *V*_*A*_ was most noticeably increased by deleterious mutations with 2*N*_*e*_*s* =1 –10 and *h* ≤ 0.2, consistent with previous work on the parameters most conducive to AOD (14).

**Figure 1.**
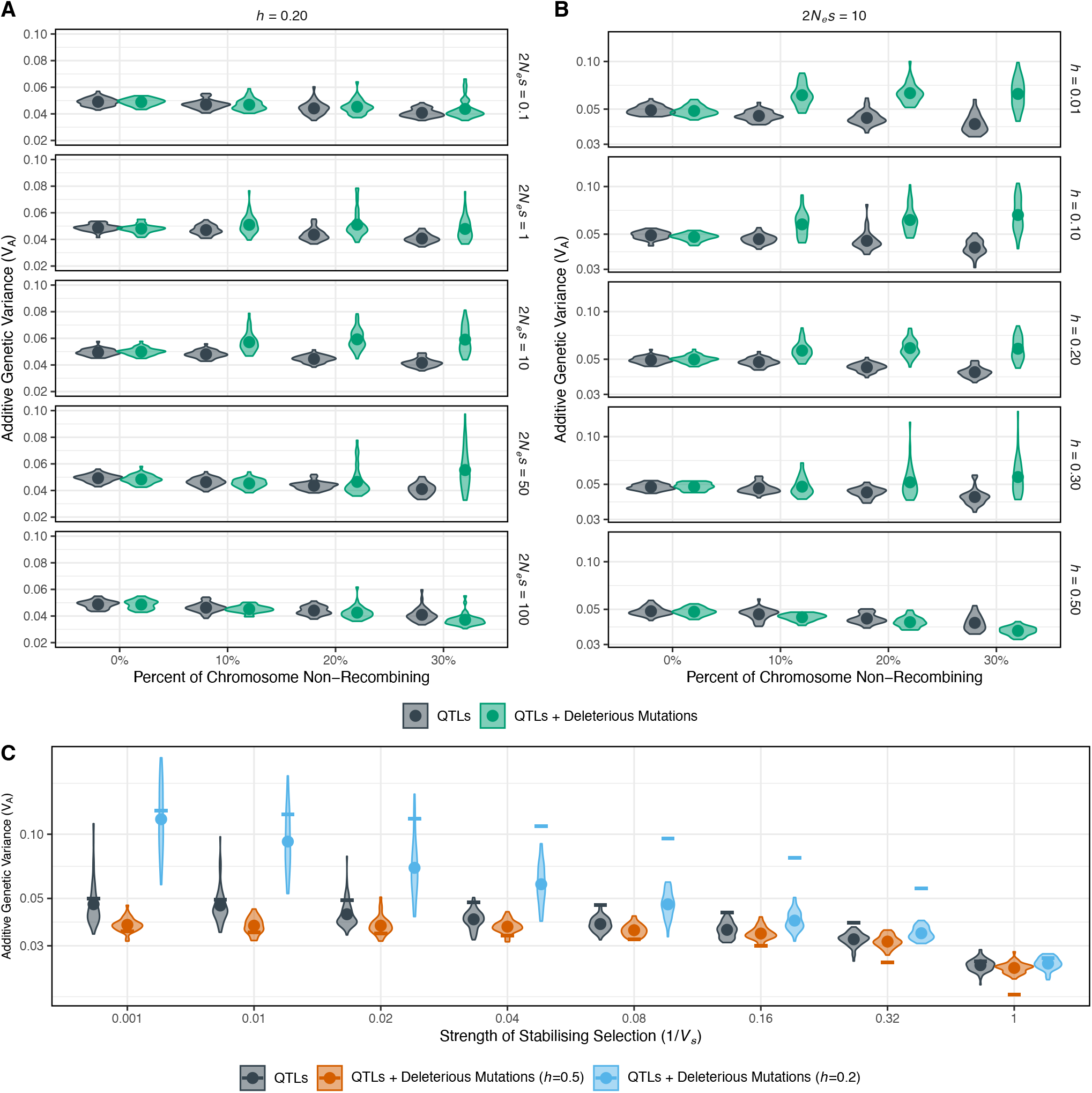
Additive genetic variance of a trait under stabilising selection with or without unconditionally deleterious mutations. **A)** *V*_*A*_ for models with or without partially recessive deleterious mutations across a range of selection strengths and genomes with an increasing fraction of non-recominbining sequence. **B**) *V*_*A*_ for models with or without mildly deleterious mutations across a range of dominance coefficients. **C)** The effect of AOD on *V*_*A*_ diminishes with an increasing strength of stabilising selection. Panel C shows simulations modelling a 30% non-recombining chromosomes with deleterious mutaitons with 2*N*_*e*_*s* = 10. Theoretical expectations (see Methods) for *V*_*A*_ in a freely recombining genome are shown as solid horizontal lines. For each parameter combination we performed 40 replicates; the distribution of values of these are shown as violins, the means are shown as solid dots.

The effect of AOD on *V*_*A*_ depends on the strength of stabilising selection on the trait. Under stabilising selection, the selection coefficients for individual QTL alleles are determined by the magnitude of their phenotypic effects and width of the fitness function for the trait (1) – see Methods. Figure 1C shows that, as the strength of stabilising selection increases, the relative effect of unconditionally deleterious mutations on *V*_*A*_ diminishes. The observed values of *V*_*A*_ in the QTL-only models were slightly smaller than theoretical expectations for the case of free recombination (see Methods), consistent with the generation of negative linkage disequilibrium among the QTLs (the Bulmer effect) (1). Li and Berg (15) recently showed that BGS caused by deleterious mutations with *h=*0.5 can further decrease *V*_*A*_ for a trait under stabilising selection, and we replicated this result across a range of selection strengths (Figure 1C and Methods). However, when partially recessive deleterious mutations were included, AOD tended to increase *V*_*A*_, especially when stabilising selection was weak (Figure 1C and Methods).

These results imply that the DFE for new deleterious mutations is, therefore, a crucially important parameter that may determine how *V*_*A*_ is maintained in populations. We therefore incorporated DFEs based on estimates for nonsynonymous mutations in three species of plants into our simulations (Figure 2A, Methods). The DFEs with the largest fraction of mildly deleterious mutations (1 ≥ 2*N*_*e*_*s* ≤ 10) exhibited the greatest *V*_*A*_ (Figure 2B). The DFEs that we modelled were inferred for nonsynonymous sites, which tend to exhibit very high levels of conservation, while non-coding DNA, which may outnumber protein coding sites, may be under weaker constraint (16, 17). Thus, when integrating across whole genomic regions, it seems likely that the DFE will contain a large fraction of mildly deleterious mutations, so that our results may underestimate the effects of linked deleterious mutations on *V*_*A*_. Nevertheless, the effects of AOD on *V*_*A*_ were evident with the DFEs we assumed here (Figure 2B).

**Figure 2.**
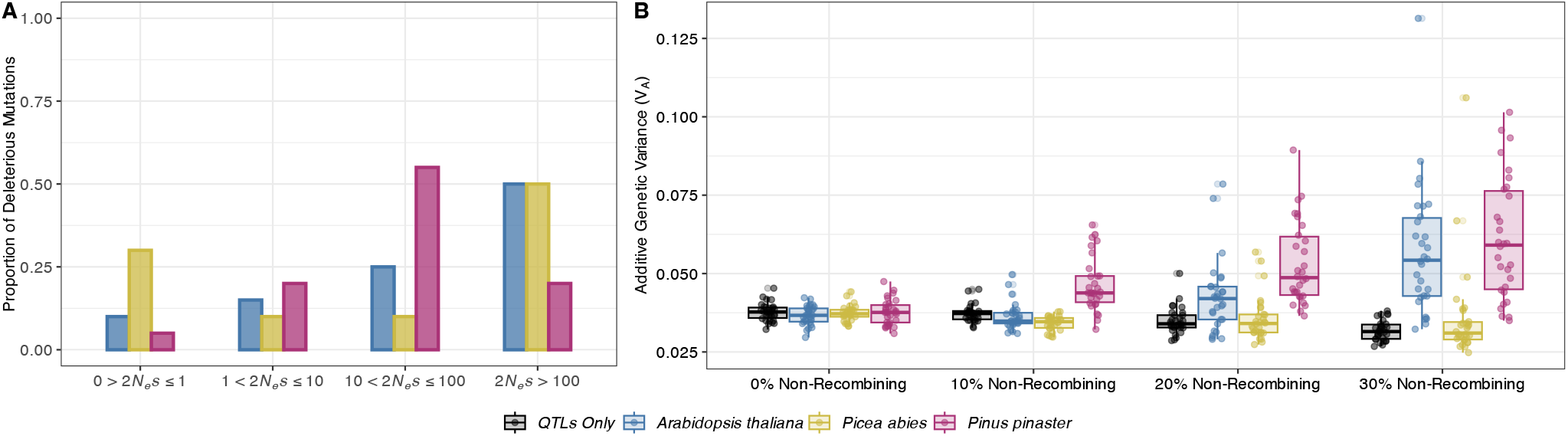
The shape of the DFE for new mutations determines the prevalence of associative overdominance and its effects on *V*_*A*_. **A)** DFEs for three plant species, represented as the proportion of new mutations falling into four ranges of selective effect. **B**) *V*_*A*_ maintained in populations experiencing deleterious mutations drawn from those DFEs in genomes with increasing proportions of non-recombining sequence. We assumed that the most strongly deleterious mutations (2*N*_*e*_*s* ≥ 25) drawn from these DFEs had *h* = 0.02 while the remaining deleterious mutations had *h* = 0.25. The quantitative trait was subject to weak stabilising selection (*V*_*s*_ = 25). For each parameter combination in b) we performed 50 replicates (points); the interquartile range and median values from these are shown as boxplots.

*V*_*A*_ may be maintained by AOD in eukaryotic genomes with deleterious mutations segregating in repulsion, so converting that *V*_*A*_ into genetic gain will come at a cost. By applying truncation selection to our simulated populations, we saw predictable responses to selection (Figure 3A) along with clear declines in the mean fitnesses of selected lines (Figure 3B). Indeed, the greatest decreases in fitness occurred in genomes with the largest non-recombining regions, and hence with the greatest contribution to *V*_*A*_ from AOD (Figure 3B). Many agriculturally important plant species’ genomes contain large regions with low frequencies of recombination (9), and signals consistent with AOD have been reported in some of these (12, 13). Selecting on traits for which *V*_*A*_ is partially maintained by AOD could help explain the trade-offs with other components of fitness that are well-documented in breeding programs (1).

**Figure 3.**
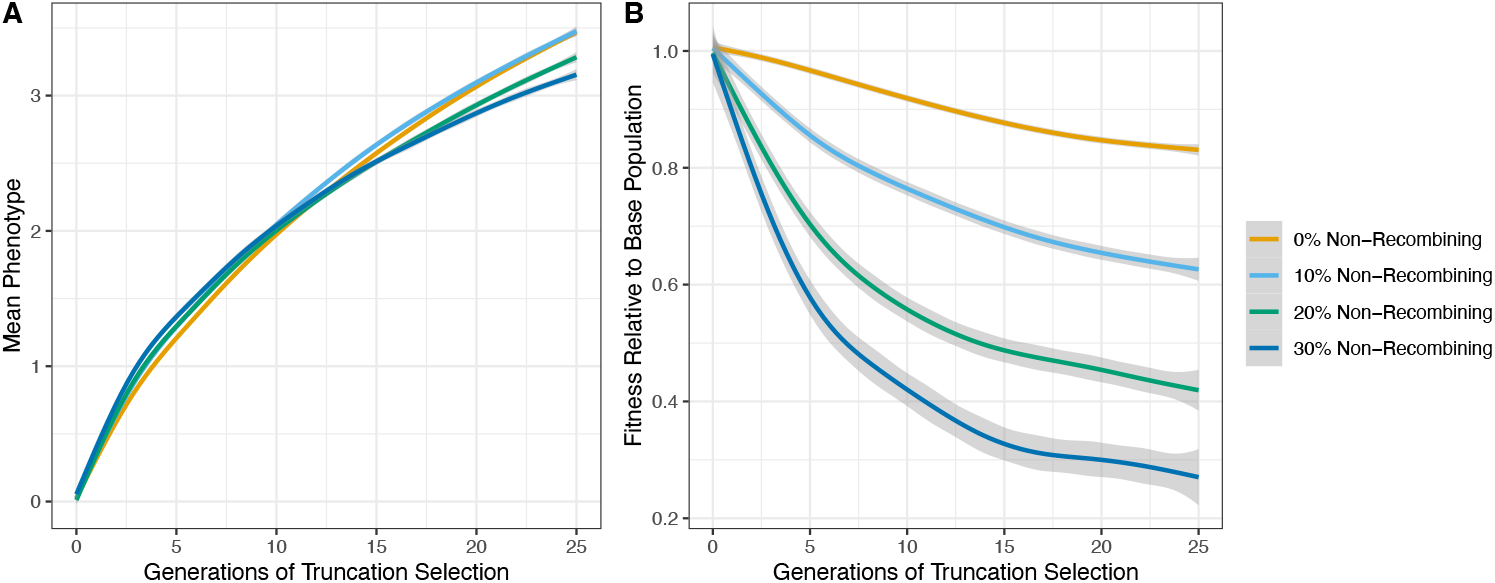
Applying truncation selection to a trait previously subject to stabilising in a population that experiences with QTLs and deleterious recessives. **A)** The response to sustained truncation selection with a 5% selection differential for 25 generations. **B)** Mean fitness of selected lines relative to the base population. The simulations assume the DFE for *Pinus pinaster*. Each line shows the average of 40 simulation replicates ± standard errors in the grey ribbons.

It seems reasonable to expect that both BGS and AOD could affect levels of *V*_*A*_ across eukaryotic species. *V*_*A*_ maintained in this way could contribute to the weak signals of stabilising selection found in natural populations (3), narrow the possible paths that evolution by natural selection can take (1). Our results, along with the recent study by Li and Berg (15), highlight the important role of selection at linked sites in affecting the maintenance of genetic variation in quantitative traits by mutation-selection balance.

## Methods

See Supplementary Text.

## Acknowledgements

Thanks to the Booker and Whitlock labs for feedback and to Peter Keightley, Loren Rieseberg, Michael Whitlock and Jarrod Hadfield for discussions. This work was supported by an NSERC Discovery grant awarded to TRB (RGPIN-2024-04434) and an NSERC Undergraduate Student Research Award awarded to YZ. PJ was supported by the NIH award R35GM154969.

## Methods

### Simulation model

We simulated a randomly mating,Wright-Fisher populations with *N* = 5,000 diploid individuals using the forward-in-time package SLiM (v 5.0; Haller et al., 2026). The simulated genome was a single 10-Mbp chromosome. All sites in the genome could potentially mutate to functional alleles. The overall mutation rate for functional alleles was *µ* = 5 × 10^−9^/bp. Each new mutation was assigned as a QTL or as unconditionally deleterious with a probability of 0.5 (i.e. the mutation rate for each mutation type was 2.5 × 10^−9^/bp). The mutation rate was selected on the assumption that approximately 5% of the genome (and hence new mutations) would be functional. If the neutral mutation rate were approximately 20 times greater, the expected genetic diversity at neutral sites would be 4*N*_*e*_*µ =* 0.002. QTL mutations were modeled as draws from a normal distribution with a mean of 0.0 and a standard deviation of 0.01. A diploid individual’s phenotype (*z*_*i*_) was the additive combination of the QTLs it possessed (i.e. the dominance coefficient was *h*=0.5). The fitness effects of deleterious mutations were either modelled as fixed effects or as draws from a DFE. Fixed selection coefficients of γ_*d*_ = 2*N*_*e*_*s*_*d*_ (*s*_*d*_ being the reduction in relative fitness for homozygotes) and a range of dominance coefficients from almost complete recessivity to additivity (0.01 ≤ *h* ≤ 0.5) were simulated. The fitness effects of the three possible genotypes for unconditionally deleterious diallelic loci were 1, 1 – *hs* and 1 – *s*.

We selected a set of three published DFEs from plant species inferred by assuming gamma distributions of γ_*d*_ : *Arabidopsis thaliana* (Chen et al., 2017), *Pinus pinaster* (James et al., 2023) and *Picea abies* (James et al., 2023) (Table S1). These DFEs were chosen as they each have rather different shapes with different fractions of mutations falling into the ranges of effect known to induce either background selection or associative overdominance (Zhao & Charlesworth, 2016). However, it should be noted that non-random mating can substantially bias estimates of the DFE from population genomic data (Daigle & Johri, 2025). For simulations assuming these DFEs we assigned a dominance coefficient of

*h* = 0.02 to deleterious mutations with a selection coefficient *s*_*d*_ < −0.002 and a dominance coefficient of *h* = 0.25 to those with a selection coefficient *s*_*d*_ > −0.002. Figure 2A in the main text shows the three DFEs, the proportion of mutations falling into different categories of mutational effects as well as the fraction of mutations with different dominance coefficients.

**Table S1.** Parameters of the DFEs for the three plant species analyzed in the main text. All DFEs are gamma distributions.

| Species | Shape Parameter | Mean $N_e s$ | Reference |
| --- | --- | --- | --- |
| *Arabidopsis thaliana* | 0.345 | -353 | Chen et al., 2017 |
| *Pinus pinaster* | 0.732 | -64.1 | James et al., 2023 |
| *Picea abies* | 0.0972 | -46700 | James et al., 2023 |

To explore the effects of recombination landscape on *V*_*A*_ for the quantitative trait, we ran each simulation across four recombination maps that varied the size of a central non-recombing region (0 - 30%, in 10% increments), while keeping the genome-wide average recombination rate constant at *c* = 2.5 × 10^−7^/bp (4*N*_*e*_*c* = 0.005). To ensure that the simulations reached selection-mutation-drift equilibrium, we implemented 30,000 (15*N*_*e*_) generations of burn-in. For each parameter combination we ran 50 simulation replicates.

Selection was modeled as the product of stabilising selection acting on the quantitative trait and the relative fitness of an individual due to unconditionally deleterious mutations. We modelled stabilising selection on the quantitative trait using the standard expression for Gaussian stabilizing selection (Walsh & Lynch, 2018) to calculate an individual’s fitness *W*_*i*_:

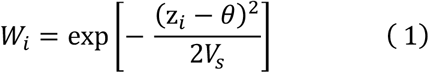

where *V*_*s*_ is the “variance” of the Gaussian fitness function, *z*_*i*_ is the phenotype of the *i*th individual, and *θ* is the phenotypic optimum, which is set to 0 in all simulations. We assumed SLiM’s default setting for unconditionally deleterious mutations (i.e. multiplicative fitnesses) to calculate an individual’s relative fitness, which was then multiplied by *W*_*i*_.

### Calculating *V*_*A*_

We used the regression of midparent mean on offspring phenotypes to compute *V*_*A*_ for the quantitative trait. Specifically, after 30,000 generations in the simulations, we recorded the parents of all offspring. We then computed the slope (*β*) of the regression of midparent mean phenotype on offspring phenotype to compute narrow sense heritability (*h*^*2*^), and converted this to *V*_*A*_.

### Artificial Selection

We modelled artificial selection on our simulated population by implementing a model of truncation selection. For 100 generations following the burn-in, individuals with quantitative trait values greater than the 90th percentile were used as parents for the next generation (i.e. a selection intensity of 10%). During this period of truncation selection, we relaxed selection on the quantitative trait but recorded each individual’s relative fitness (distinct from the effect of truncation selection) from the unconditionally deleterious mutations they possessed.

### Expected *V*_*A*_ under stabilising selection

We used the method of McVean & Charlesworth (1999) find an expression for the nucleotide site diversity (heterozygosity) at sites under stabilising selection, using the infinite sites assumption that new mutations arise only at non-segregating sites, corresponding to a negligible proportion of segregating sites. On the assumption of no directionality to mutations, so that mutations increasing and decreasing the trait value both occur at rate *m* per generation, the symmetry of the model implies that the probability of fixation of a mutation with average effect *a* is the same as one with average effect – *a* . Similarly, the expected total heterozygosity *H* contributed by an A_1_ mutation on its way to fixation or loss is the same as that for an A_2_ mutation. To obtain the net nucleotide site diversity, *π, H* is multiplied by the rate of input of new mutations, *N*_*H*_*µ*, yielding:

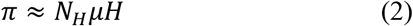

where *N*_*H*_ is the number of haploid copies of a locus (2*N* in the case of a diploid, autosomal locus).

*H* is given by integrals of somewhat complicated forms that depend on the variance effective population size, *N*_*e*_, and the specific model of selection (Ewens, 2004, Chapter 4; (Charlesworth, 2022). However, for sufficiently weak selection, useful approximations can be found (Charlesworth, 2022). For a wide class of selection models, the rate of change of allele frequency *x* in units of 2*N*_*e*_ generations can be written as:

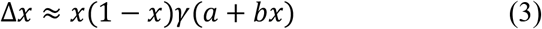

where *γ* is the scaled selection coefficient, 2*N*_*e*_*s*, and *a* and *b* are constants that represent a linear function of *x*.

In the case of sufficiently weak stabilizing selection, the nor-optimal selection model can be approximated by the quadratic deviations model of Wright (1935) (see Bürger 2000, p.201), and we have the following simple relations:

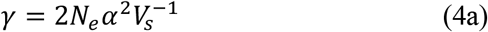

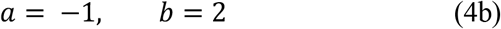

Equation (7) of Charlesworth (2022) gives the following second-order approximation with respect to terms in *g* :

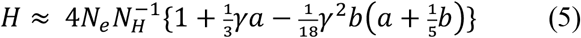

Using Equation (1), we obtain‘;

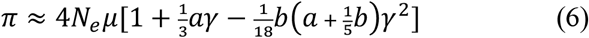

Substituting *a* = – 1 and *b* = 2 for the case of stabilizing selection, this becomes:

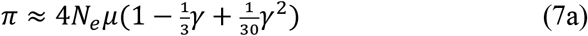

Writing *S* = 2*N*_*e*_/*V*_*s*_, we obtain:

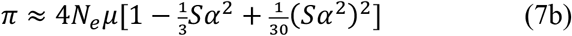

The corresponding expression for the additive variance contributed by a single site is:

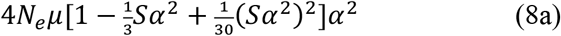

Assuming normality of the distribution of *a*, ignoring linkage disequilibrium, and using the fact that the fourth and sixth moments of the normal distribution are 3*σ*^4^ and 15*σ*^6^, respectively. the approximate expected additive variance contributed by *n* additive loci with expected mean squared average effect *σ*^2^is given by:

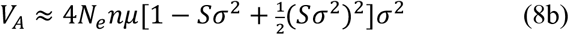

The corresponding ratio of *V*_*A*_ to its value in the absence of stabilizing selection is given by:

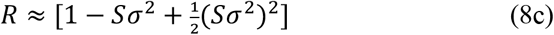

Equations (7) are likely to be inaccurate for values of *Sσ*^2^ of the order of 1 or more, due to contributions from higher-order terms in *Sσ*^2^, so that numerical evaluations of the relevant integrals must be used, as described in (Charlesworth, 2022).

This approach can be extended to situations where deleterious mutations are segregating in the background, causing background selection (BGS) and/or associative overdominance (AOD). When *h* = 0.5, only BGS will occur. Its approximate effect can be found as follows for the case when the central proportion of the chromosome is non-recombining and the rest is recombining at a rate that would contribute a total map length of *M* in Morgans to the chromosome if there were no region without recombination: *M* = *mr*, where *m* is the total number of nucleotide sites on the chromosome, and *r* is the rate of crossing over per site. The total mutation rate to deleterious alleles per haploid genome is denoted by *U*.

Let *x* be the proportion of the chromosome occupied by the central region. For this region, the ratio of neutral diversity in the presence of BGS to that in its absence (*B*_*c*_) is given approximately by the formula of (Charlesworth et al., 1993):

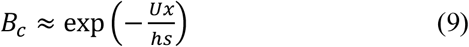

This implies that the effective population size for sites located in the central region is *B*_*c*_*N*_*e*_. The corresponding additive variance for the quantitative trait for a given strength of stabilizing selection *V*_*s*_ is *V*_*Ac*_ = *V*_*A*_(*B*_*c*_*N*_*e*_, *V*_*s*_), which can be found from the above approach by substituting *B*_*c*_*N*_*e*_ for *N*_*e*_ into either the approximate or exact formulae.

For the distal remainder of the chromosome, the following approximate formula for the relative neutral diversity under BGS can be used (Hudson & Kaplan, 1995; Nordborg et al., 1996):

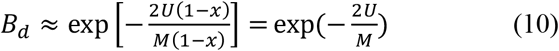

The additive variance contributed by the distal portion of the chromosome can be found by using an effective population size *B*_*c*_*N*_*e*_ in the expressions for *V*_*A*_, such that *V*_*Ad*_ = *V*_*A*_(*B*_*d*_*N*_*e*_, *V*_*s*_). The overall additive variance with BGS is thus given by:

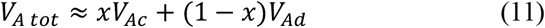

In the case shown in Figure (1c), *x* = 0.3, *s* = 0.001, *U* = 0.5 x 10^7^ x 5 x 10^−9^ = 0.025. With *h* = 0.5, *Ux*/*hs* = 15, so that *B*_*c*_ ≈ 3.03 x 10^−7^ ≈ 0. There is thus no contribution to *V*_*A tot*_ from the central region. For the distal regions, the map length of the chromosome if there were no central region is *M* = 10^7^ x 2 x 10^−7^ = 2, so that 2*U*/*M* = 0.025, giving *B*_*d*_ ≈ 0.0975. For practical purposes this can equated to 1, so that *V*_*Ad*_ is given by *V*_*A*_(*N*_*e*_, *V*_*s*_), the corresponding value of *V*_*A*_ in the absence of BGS, and *V*_*A tot*_ is simply (1 − *x*)*V*_*A*_(*N*_*e*_, *V*_*s*_).

With *h* = 0.2 and a large central non-recombining regions, the simulations show that AOD rather than BGS is operating in the central region. There is currently no theoretical prediction for this situation, but the following *ad hoc* method can be used. Assume that the central region alone is subject to AOD, whereas the recombining distal regions obey Equation (9) for the effects of BGS alone. Assume also that the simulation results for the weakest intensity of stabilizing selection plus deleterious mutations are a good proxy for the case of the effects on neutral diversity of deleterious mutations alone. Let the ratio of *V*_*A*_ for this case to *V*_*A*_ in the absence of deleterious mutations be *R*, which can be estimated from the simulation results for 1/*V*_*s*_ = 0.001, and let the ratio for the central region alone be *A*_*c*_. We then have the following relation:

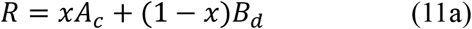

so that

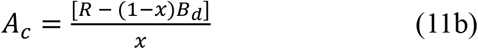

Assume also that *V*_*A*_ for the distal regions is approximately the same for *h* = 0.5, since there is no explicit dependence on *hs* in Equation (9). By analogy with the derivation for the case of BGS alone, *V*_*A*_ for the central region for a given strength of stabilizing selection is obtained by replacing *N*_*e*_ with *A*_*c*_*N*_*e*,_, giving 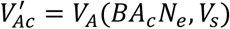. The overall value of *V*_*A*_ is then given by:

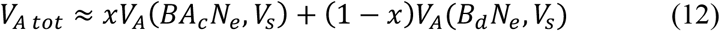

For the case shown in Figure (1c), the simulations give an estimate of *R* = 2.578 with 1/*V*_*s*_ = 0.001. Using the above results for BGS alone, *B*_*d*_ for this case is approximately 1. The approximate and exact theoretical values of 1/*V*_*s*_ = 0.001 for this case are both 0.05 (see Table S2). Equation (11a) thus gives *A*_*c*_ = (2.58 – 0.7)/0.3 = 6.27.

Table S2 shows the results of the relevant calculations compared with the simulation results.

Overall, the exact theoretical predictions for *V*_*A*_ in the absence of deleterious mutations are fairly close to the simulation values until 1/*V*_*s*_ = 20, and the approximate predictions are close to the exact values up to 1/*V*_*s*_ = 0.32. The predicted values are large than the simulated values, probably reflecting the effect of the negative linkage disequilibrium generated by stabilizing selection in reducing *V*_*A*_ below the value obtained by summing the effects of individual loci: the “Bulmer effect” (Bulmer, 1971, 1974). Somewhat surprisingly, similarly good agreement is found with BGS alone (*h* = 0.5), but with *V*_*A*_ mostly being underestimated by the exact theoretical predictions, and a very poor performance of the approximate predictions for 1/*V*_*s*_ > 0.32. The underestimation probably results from the deterministic model of mutation and selection used to obtain Equation (9) not being completely accurate for a finite population (Nordborg et al., 1996).

The agreement between theory and simulations for the case of AOD together with BGS (*h* = 0.2) is substantially worse, which is not surprising in view of the heuristic nature of the approach.

Nevertheless, it is quite good for 1/ *V*_*s*_ up to 0.04 and then for 1/*V*_*s*_ ≥ 1.

Approx. and Exact refer to the use of Equation (7c) for the effect of stabilizing selection versus the use of numerical evaluation of the integrals for *H*, as described in Charlesworth (2022).

**Table S2.** Theoretical predictions of the additive variance for the parameters used in Figure 1C, compared with the means from the simulations.

**No deleterious mutations**
| $1/V_s$ | Approx. | Exact | Simulation |
| --- | --- | --- | --- |
| 0.001 | 0.05 | 0.05 | 0.0485 |
| 0.01 | 0.0495 | 0.0494 | 0.0472 |
| 0.02 | 0.049 | 0.049 | 0.0426 |
| 0.04 | 0.048 | 0.0479 | 0.0401 |
| 0.08 | 0.046 | 0.0465 | 0.038 |
| 0.16 | 0.043 | 0.0429 | 0.0358 |
| 0.32 | 0.038 | 0.0385 | 0.0323 |
| 1 | 0.0389 | 0.0253 | 0.0244 |
| 2 | 0.106 | 0.0168 | 0.0188 |
| 20 | 14.6 | 0.0126 | 0.0042 |

**Deleterious mutations with $h = 0.5$**
| $1/V_s$ | Approx. | Exact | Simulation |
| --- | --- | --- | --- |
| 0.001 | 0.035 | 0.035 | 0.0377 |
| 0.01 | 0.0346 | 0.0346 | 0.0374 |
| 0.02 | 0.0343 | 0.0343 | 0.0375 |
| 0.04 | 0.0334 | 0.0334 | 0.037 |
| 0.08 | 0.0322 | 0.0322 | 0.0356 |
| 0.16 | 0.0301 | 0.03 | 0.0343 |
| 0.32 | 0.0266 | 0.0251 | 0.0315 |
| 1 | 0.0272 | 0.0177 | 0.0237 |
| 2 | 0.0742 | 0.0118 | 0.0184 |
| 20 | 1.48 | 0.0088 | 0.0041 |

**Deleterious mutations with $h = 0.2$**
| $1/V_s$ | Approx. | Exact | Simulation |
| --- | --- | --- | --- |
| 0.001 | 0.129 | 0.129 | 0.125 |
| 0.01 | 0.124 | 0.124 | 0.0973 |
| 0.02 | 0.118 | 0.118 | 0.0732 |
| 0.04 | 0.109 | 0.109 | 0.0596 |
| 0.08 | 0.0975 | 0.0955 | 0.0474 |
| 0.16 | 0.104 | 0.0776 | 0.0396 |
| 0.32 | 0.228 | 0.0556 | 0.0344 |
| 1 | 2.89 | 0.0263 | 0.0248 |
| 2 | 12.5 | 0.0151 | 0.0187 |
| 20 | 1352 | 0.001 | 0.0041 |

## Notes

### Competing Interest Statement

The authors have declared no competing interest.

## References

1. B. Walsh, M. Lynch, Evolution and selection of quantitative traits (Oxford University Press, 2018).

2. L. C. Zijmers, K. L. Abson, J. D. Hadfield, A. Eyre-Walker, Levels of additive genetic variation vary substantially between species. PLoS Biol. 24, e3003819 (2026).

3. T. Johnson, N. Barton, Theoretical models of selection and mutation on quantitative traits. Phil. Trans. R. Soc. B 360, 1411–1425 (2005).

4. J. G. Kingsolver, et al., The strength of phenotypic selection in natural populations. Am. Nat. 157, 245–261 (2001).

5. Y. B. Simons, et al., Simple scaling laws control the genetic architectures of human complex traits. PLoS Biol. 23, e3003402 (2025).

6. X.-S. Zhang, W. G. Hill, Genetic variability under mutation selection balance. Trends Ecol. Evol. 20, 468–470 (2005).

7. N. H. Barton, A. M. Etheridge, A. Véber, The infinitesimal model: Definition, derivation, and implications. Theor. Popul. Biol. 118, 50–73 (2017).

8. J. F. Crow, Mutation, mean fitness, and genetic load. Oxford Series in Evolutionary Biology 9, 3–42 (1993).

9. T. Brazier, S. Glémin, Diversity and determinants of recombination landscapes in flowering plants. PLoS Genet. 18, e1010141 (2022).

10. B. Charlesworth, J. D. Jensen, Effects of selection at linked sites on patterns of genetic variability. Annu. Rev. Ecol. Evol. Syst. 52, 177–197 (2021).

11. D. M. Waller, Addressing Darwin’s dilemma: Can pseudo-overdominance explain persistent inbreeding depression and load? Evolution 75, 779–793 (2021).

12. E. Rodgers-Melnick, et al., Recombination in diverse maize is stable, predictable, and associated with genetic load. Proc. Natl. Acad. Sci. U. S. A. 112, 3823–3828 (2015).

13. M. Salson, et al., Interplay between large low-recombining regions and pseudo-overdominance in a plant genome. Nat. Commun. 16, 6458 (2025).

14. L. Zhao, B. Charlesworth, Resolving the conflict between associative overdominance and background selection. Genetics 203, 1315–1334 (2016).

15. X. Li, J. J. Berg, Background selection in recombining genomes and its consequences for the maintenance of variation in complex traits. Proc. Natl. Acad. Sci. U. S. A. 123, e2513613123 (2026).

16. J. L. Campos, L. Zhao, B. Charlesworth, Estimating the parameters of background selection and selective sweeps in Drosophila in the presence of gene conversion. Proc. Natl. Acad. Sci. U. S. A. 114, E4762–E4771 (2017).

17. D. L. Halligan, et al., Contributions of protein-coding and regulatory change to adaptive molecular evolution in murid rodents. PLoS Genet. 9, e1003995 (2013).

## Supplementary References

Bulmer, M. G. (1971). The effect of selection on genetic variability. The American Naturalist, 105(943), 201–211.

Bulmer, M. G. (1974). Linkage disequilibrium and genetic variability. Genetical Research, 23(3), 281–289.

Bürger, R. (2000). The mathematical theory of selection, recombination, and mutation. John Wiley & Sons.

Charlesworth, B. (2022). The effects of weak selection on neutral diversity at linked sites. Genetics, 221(1), iyac027.

Charlesworth, B., Morgan, M. T., & Charlesworth, D. (1993). The effect of deleterious mutations on neutral molecular variation. Genetics, 134(4), 1289–1303.

Chen, J., Glémin, S., & Lascoux, M. (2017). Genetic diversity and the efficacy of purifying selection across plant and animal species. Molecular Biology and Evolution, 34(6), 1417–1428.

Daigle, A., & Johri, P. (2025). Hill-Robertson interference may bias the inference of fitness effects of new mutations in highly selfing species. Evolution; International Journal of Organic Evolution, 79(3), 342–363.

Ewens, W. J. (2004). Mathematical population genetics 1: Theoretical introduction (2nd ed.) [PDF]. Springer.

Haller, B. C., Ralph, P. L., & Messer, P. W. (2026). SLiM 5: Eco-evolutionary simulations across multiple chromosomes and full genomes. Molecular Biology and Evolution, 43(1). 10.1093/molbev/msaf313

Hudson, R. R., & Kaplan, N. L. (1995). Deleterious background selection with recombination. Genetics, 141(4), 1605–1617.

James, J., Kastally, C., Budde, K. B., González-Martínez, S. C., Milesi, P., Pyhäjärvi, T., Lascoux, M., & GenTree Consortium. (2023). Between but not within-species variation in the distribution of fitness effects. Molecular Biology and Evolution, 40(11), msad228.

McVean, G. A. T., & Charlesworth, B. (1999). A population genetic model for the evolution of synonymous codon usage: patterns and predictions. Genetics Research, 74(2), 145–158.

Nordborg, M., Charlesworth, B., & Charlesworth, D. (1996). The effect of recombination on background selection. Genetical Research, 67(2), 159–174.

Walsh, B., & Lynch, M. (2018). Evolution and selection of quantitative traits. Oxford University Press.

Wright, S. (1935). The analysis of variance and the correlations between relatives with respect to deviations from an optimum. Journal of Genetics, 30(2), 243–256.

Zhao, L., & Charlesworth, B. (2016). Resolving the conflict between associative overdominance and background selection. Genetics, 203(3), 1315–1334.

